# Salubrinal induces fetal hemoglobin expression via the stress-signaling pathway in human sickle erythroid progenitors and sickle cell disease mice

**DOI:** 10.1101/2021.12.13.472406

**Authors:** Nicole H. Lopez, Biaoru Li, Chithra Palani, Umapathy Siddaramappa, Mayuko Takezaki, Hongyan Xu, Wenbo Zhi, Betty S. Pace

## Abstract

Sickle cell disease (SCD) is an inherited blood disorder caused by a mutation in the *HBB* gene leading to hemoglobin S production and polymerization under hypoxia conditions leading to vaso-occlusion, chronic hemolysis, and progressive organ damage. This disease affects ∼100,000 people in the United States and millions worldwide. An effective therapy for SCD is fetal hemoglobin (HbF) induction by pharmacologic agents such as hydroxyurea, the only Food and Drug Administration-approved drug for this purpose. Therefore, the goal of our study was to determine whether salubrinal (SAL), a selective protein phosphatase 1 inhibitor, induces HbF expression through the stress-signaling pathway by activation of p-eIF2α and ATF4 trans-activation in the γ-globin gene promoter. Sickle erythroid progenitors treated with 24µM SAL increased F-cells levels 1.4-fold (p=0.021) and produced an 80% decrease in reactive oxygen species. Western blot analysis showed SAL enhanced HbF protein by 1.6-fold (p=0.0441), along with dose-dependent increases of p-eIF2α and ATF4 levels. Subsequent treatment of SCD mice by a single intraperitoneal injection of SAL (5mg/kg) produced peak plasma concentrations at 6 hours. Chronic treatments of SCD mice with SAL mediated a 2.3-fold increase in F-cells (p=0.0013) and decreased sickle erythrocytes supporting *in vivo* HbF induction.

## INTRODUCTION

Sickle Cell Disease (SCD) is a group of inherited blood disorders caused by point mutations in the *HBB* gene on chromosome 11. About 100,000 African Americans are affected with SCD in the United States, while millions suffer worldwide. The clinical symptoms of SCD include chronic hemolytic anemia, recurrent acute pain, and progressive organ damage. However, higher fetal hemoglobin (HbF; α_2_γ_2_) levels ameliorate clinical severity in people with SCD due to inhibition of hemoglobin S (HbS) polymerization in erythrocytes. Data generated in the Cooperative Study of Sickle Cell Disease, demonstrated that HbF >8.6% correlated with improved long-term survival in adults with SCD (1). Therefore, reactivation of *HBG* gene expression by pharmacologic agents is an effective therapeutic strategy for treating SCD (2).

Several drugs induce HbF *in vivo* including hydroxyurea (HU), arginine butyrate, and decitabine, among others; however, undesirable side effects have hampered clinical development of some. Hydroxyurea is a short-acting cytotoxic ribonucleotide reductase inhibitor, which increases nitric oxide levels and activation of soluble guanylyl cyclase leading to *HBG* activation (3). Arginine Butyrate is a histone deacetylase inhibitor that rapidly stimulates HbF expression via mRNA stabilization (4). By contrast, decitabine is a DNA methyltransferase inhibitor mediating hypomethylation of the *HBG* proximal promoters and gene activation (5). Currently, HU is the only Food and Drug Administration (FDA)-approved drug to induce HbF synthesis in SCD patients. However, only two-third of adults respond to HU treatment and suffer dose-limiting side effects, such as bone marrow suppression, making this agent less than ideal(3). After two decades of clinical investigation, three new drugs were recently approved by the FDA for the treatment of pain and anemia in SCD including L-glutamine (6), voxelotor (7) and crizanlizumab (8), however, these agents are not HbF inducers. Therefore, we sought to develop a novel, HbF inducing agents with the potential to reduce oxidative stress in SCD erythrocytes.

Salubrinal (SAL) is a semi-permeable synthetic compound and selective inhibitor of protein phosphatase 1 (PP1). The net result of this action is to interfere with the recruitment of phosphorylated eukaryotic initiation factor 2α (p-eIF2α) to PP1 which increases p-eIF2α levels to activate downstream targets such as ATF4 (activating transcription factor 4) (9). Furthermore, Chen, et al demonstrated that SAL rescues anemia in the DDRGK1^F/F^ knockout mouse model (10). These animals have hematopoietic dysfunction including death of hematopoietic stem and progenitor cells and acute anemia. More directly related to *HBG* gene regulation, Hahn et al (2) demonstrated in normal erythroid progenitors, that SAL prevents dephosphorylation of p-eIF2α, thus mediating ATF4 activation to regulate the integrated stress response pathway. In addition, SAL increases HbF production by a posttranscriptional mechanism, without affecting cell proliferation, differentiation, or *HBG* mRNA levels.

To expand these *in vitro* studies, we initially treated K562 cells with SAL mediating *HBG* activation with a simultaneous increase in HbF, p-eIF2α and ATF-4 protein levels. To investigate the role of ATF4 in the mechanism of HbF induction by SAL, we identified predicted ATF4 binding sites at -835bp upstream of *HBG2*, and the second intron of *HBB* by ChIPseq analysis reported in the ENCODE (encyclopedia of DNA elements) database. Our confirmatory ChIP assay demonstrated *in vivo* ATF4 binding at the -835*HBG2* motif and to a lesser degree for *HBB*. Studies with siATF4 and SAL treatment support in part HbF induction by ATF4. Therefore, to expand the field we investigated sickle erythroid progenitors generated from peripheral blood mononuclear cells (PBMC). SAL mediated a significant increase in HbF positive cells (F-cells), the mean fluorescent intensity (MFI) a measure to quantify HbF levels per cell and HbF protein levels by Western blot, along with a significant decrease in reactive oxygen species (ROS). Moreover, SAL increased p-eIF2α and ATF4 expression in SCD progenitors. Subsequent treatment of the Townes SCD transgenic mice for four weeks with SAL produced a significant increase in F-cells with minimal hematopoietic toxicity. Finally, the percentage of sickled erythrocytes decreased by 66% at four weeks supporting a change in peripheral blood phenotype. The novel observation of HbF induction in SCD transgenic mice support pre-clinical development of SAL as a treatment option for SCD.

## MATERIALS AND METHODS

### Drug treatments

Salubrinal (SML0951) purchased from Sigma-Aldrich (St. Louis, MO) was reconstituted in 100% dimethyl sulfoxide (DMSO; BP231) for a 50mM stock solution. A second SAL (SC202332) reagent was purchased from Santa Cruz Biotechnology (Dallas, TX) and reconstituted in water at a 1mM concentration stock. SAL dissolved in water was added to the cells containing media at the corresponding concentrations and did not precipitate out of solution at any point in the 48-hour incubation. Hemin (51280), hydroxyurea (H8627) and L-cysteine (168149) were purchased from Sigma-Aldrich. Hemin and hydroxyurea are known HbF inducers and were used as controls. Cysteine is a negative control since it does not induce HbF synthesis.

### Tissue Culture

K562 cells cultured in Iscove’s Modified Dulbecco medium, fetal bovine serum (10%), penicillin and streptomycin underwent drug inductions for 48 hours, and cell viability assessed by 0.4% Trypan blue exclusion. For primary cultures, human erythroid progenitors were generated from peripheral blood mononuclear cells (PBMCs) isolated from SCD patients as previously published (11). Treatments with SAL, DMSO, cysteine, hemin, HU, and combined SAL/HU occurred on day 8 erythroid progenitors and then harvested on day 10.

### Flow Cytometry

K562 cells and erythroid progenitors were fixed with 4% paraformaldehyde, permeabilized and stained with fluorescein isothiocyanate (FITC) anti-HbF antibody (AB-19365, Abcam, Cambridge MA) and FITC-isotype IgG antibody (AB-37406). Flow cytometry was conducted on the FACSCanto (BD Biosciences, San Jose CA) as previously published (12). To quantify ROS levels, cells were incubated with 2’,7’-dichlorodihydrofluorescein diacetate (DCF-DA) (10µM; Invitrogen, Carlsbad CA) 4 hours before harvest. To determine the effect of SAL on maturation, sickle erythroid progenitors (5×10^5^) were harvested after drug inductions, fixed with 4% paraformaldehyde and then permeabilized in ice-cold acetone/methanol (4:1). The cells were stained with human FITC conjugated transferrin receptor (CD71; ThermoFisher: 11-0719-42) and human FITC conjugated glycophorin A (CD235a; ThermoFisher: 11-9886-42) antibodies and analyzed by flow cytometry.

### Western Blot

Western blot was performed as previously published (13) with antibodies purchased from Santa Cruz Biotechnology including HbF (SC-21756), HbS (SC-21757), eIF2α (SC-11386) and Tubulin (SC-53646). β-actin (AM4302) was purchased from Invitrogen (Waltham, MA) and phospho-eIF2α (Ser51) (CS-3398S) and ATF4 (CS-11815S) from Cell Signaling Technology (Danvers, MA).

### Reverse transcription-quantitative PCR (RT-qPCR) analysis

The RT-qPCR assays were performed as previously published (14). Gene-specific primers were used to quantify mRNA levels of *HBG, HBB, ATF4, BCL11A* and internal control glyceraldehyde-3-phosphate dehydrogenase (*GAPDH*); all mRNA levels were normalized to GAPDH levels before statistical analysis. ATF4 gene silencing was achieved using ATF4 siRNA (SC-35112, Santa Cruz Biotechnology) (200 nM) to transfect K562 cells for 24 h using the Dharmafect 1 system; scrambled siRNA (sc-37007, Santa Cruz Biotechnology) was used as control. After transfections, K562 cells were treated with SAL (24μM) in triplicates for 48 hours and analyzed by RT-qPCR.

### Chromatin Immunoprecipitation Assay (ChIP)

ChIP assay was performed using the Active Motif ChIP-IT High Sensitivity kit (Carlsbad, CA) per the manufacturer’s instruction. Briefly, DNA was cross-linked with 1% formaldehyde, and sonicated to shear DNA to ∼500 bp. Immunoprecipitations were performed using ATF4 antibody (SC-390063), mouse IgG and anti-RNA polymerase II. Chromatin enrichment was quantified by qPCR using primer pairs designed with Primer-BLAST software; ATF4 sites at -835*HBG2* (forward AGAATGGGGCACAGTGGATG, reverse CCAGCATCCCGACCATGATT) and the second intron *HBB* (forward GGCCCACAAGTATCACTAAGC, reverse CACTGACCTCCCACATTCCC). Additional primers included: 1) locus control region DNase I hypersensitive site 2 (LCR-HS2) (forward CCTTCTGGCTCAAGCACAGC, reverse ATAGGAGTCATCACTCTAGGC) and 2) *HBG2* cAMP response element (G-CRE) (forward CGGCGGCTGGCTAGGGATGAA, reverse CTGTGAAATGACCCATGGCG).

### Pharmacokinetics protocol

SCD mice received a single intraperitoneal (IP) injection of SAL 5mg/kg and blood was drawn at 15 min, 30 min, and 1, 2, 3 6 and 24 hours for liquid chromatography-multiple reaction monitoring mass spectrometry (LC-MRM MS) at Augusta University. The optimal collision energy and retention factor lens were determined using standards and transitions established for SAL was 481/187, 481/129 and 481/189. The integrated peak areas for transitions were calculated (Skyline software, version 20.0).

### Townes transgenic mouse treatment

The Townes SCD mouse completes hemoglobin switching from human γ-globin to β^S^-globin shortly after birth (15).. These mice display the same clinical symptoms as human SCD patients including erythrocyte sickling, hemolysis and anemia, splenomegaly, and vaso-occlusive pain. Before starting chronic treatments, mice were injected with a single dose of SAL 5 mg/kg and later sacrificed to confirm SAL did not precipitate in the peritoneum. Drug treatments in SCD mice (4-6 months old) included: 1) water control (vehicle), 2) HU 100 mg/kg/day (positive control), 3) SAL 3 mg/kg/day, and 4) SAL 5 mg/kg/day, dissolved in water. Intraperitoxsneal injections occurred 5 days a week for 4 weeks, for two independent replicates (sample size 10 mice per group). Blood collected in EDTA tubes by tail bleed at week 0, 2, and 4 was analyzed for complete blood count and differential (Micros 60 machine; HORIBA Medical/ABX Diagnostics) and F-cells and MFI quantified by flow cytometry as previously published (31). Blood stained with BD retic-count reagent (acridine orange) quantifies reticulocytes by flow cytometry.

### Giemsa Staining

Erythroid progenitors generated from sickle peripheral blood mononuclear cells and SCD mouse blood (10uL) were collected for blood smears and stained with the Sure-Stain Wright-Giemsa (CS434D; Fisher Scientific). The number of sickled erythrocytes per high power field were counted for 500 cells in triplicate per treatment condition for the mouse studies.

### Statistical Analysis

Tissue culture experiments repeated for 3-5 independent studies with three replicates per run. The unpaired student’s *t*-test was performed and *p<0.05 was considered statistically significant. The data are plotted as the mean ± standard error of the mean (SEM). For the SCD mouse studies one-sample paired student’s t-tests were performed to determine differences in blood counts, reticulocytes, F-cells, and MFI at week 0 compared to week 2 and 4 within groups (n=10). To compare the effect across treatment groups, pairwise comparisons analysis of variance (ANOVA) was performed. A level of p≤0.05 was used for statistical significance.

## RESULTS

### SAL induces HbF expression through the integrated stress response pathway

To gain insight into mechanisms of HbF induction by SAL, studies were conducted in K562 erythroleukemia cells that carries an embryonic/fetal phenotype and expresses endogenous ε-globin and γ-globin (16). After treatment with SAL and other control agents for 48 hours, RT-qPCR was performed to determine *HBG* mRNA levels normalized to *GADPH*. A significant 3.8-fold (p=0.002) and 2.8-fold (p=0.008) increase in *HBG* mRNA was observed at SAL 12µM and 18µM respectively (Fig. 1A) compared to DMSO vehicle control. As expected, hemin (Hem) produced a significant 6.2-fold increase (p=0.00013) in *HBG* synthesis. Subsequent flow cytometry with anti-FITC HbF antibody was conducted. Shown in Fig. 1B and 1C are representative histograms and quantitative F-cell levels respectively compared to untreated (UT) K562 cells; note, SAL 12µM and 18µM increased F-Cells by 1.3-fold (p=0.003) and 1.4-fold (p=0.004) respectively.

**Figure 1.**
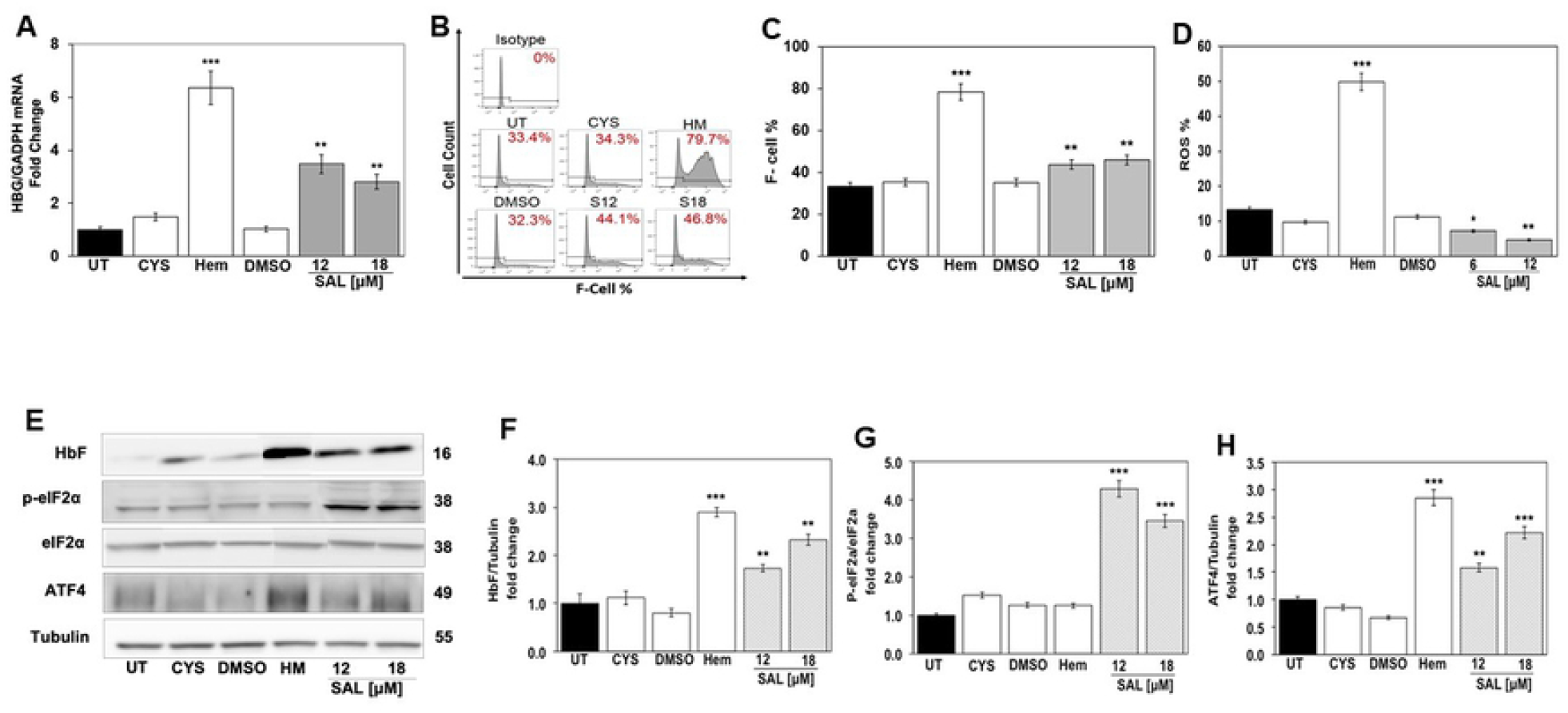
SAL increased *HBG* transcription and HbF synthesis. K562 cells were grown without treatment (UT) or treated with Cysteine 10µ M (CYS; negative control), hemin 50µ M (Hem; positive control) or salubrinal (SAL) 12μM and 18μM and vehicle control dimethyl sulfoxide (DMSO). Total RNA was isolated and whole cell lysates generated for RT-qPCR and Western blot analysis (See Material and Methods). All data are shown as the mean ± SEM (n=6) and *p<0.05; **p<0.01; *** p<0.001 was considered statistically significant. **A)** Shown in the bar graph is the *HBG/GAPDH* normalized mRNA levels generated by RT-qPCR analysis for each treatment conditions. **B)** Flow cytometry shows histograms of K562 cells under the different conditions were stained with FITC-conjugated anti-HbF antibody. **C)** Shown in bar graph is the percentage of F-cells (F-cell %)-using FACS Diva software. **D)** Shown in the bar graph is the ROS % in the different conditions after staining with DCFDA (See Materials and Methods). **E)** Western blot analysis determined HbF, p-eIF2α and ATF4 protein levels compared with tubulin (internal loading control); shown is a representative blot. Quantitative data generated by densitometry analysis quantified the expression levels of **F)** HbF normalized to tubulin, **G)** p-eIF2α normalized to total eIF2α, and **H)** ATF4 normalized to tubulin.

To measure oxidative stress, K562 cells were treated with DCF-DA (10 µM) and analyzed by flow cytometry which, showed a decrease in ROS by 50% (p=0.025) and 60% (p=0.008) at SAL 6µM and 12µM compared to 5-fold (p=0.0001) increase by hemin, known to induce oxidative stress (Fig. 1D). To gain insight into the role of stress signaling in HbF induction by SAL, Western blot (Fig. 1E) confirmed the ability of SAL to induce HbF by 2.5-fold (p=0.005) and increase p-eIF2α by 3.8-fold (p=0.00015) and ATF4 by 2.0-fold (p=0.00062) (Fig. 1F-1H). Interestingly, hemin induced ATF4 by 2.9-fold (p=0.00045) related to its ability to generate high levels of ROS in cells (Fig. 1D).

To determine how quickly SAL modulated γ-globin transcription we performed time course studies. K562 cells treated with cysteine 10µM (CYS), Hemin 50µM (Hem), DMSO, SAL 12 μM, and SAL/Hem (SAL12μM/HM50 μM) were harvested at 6 and 24 hours. Subsequent RT-qPCR at 6 hours showed no change in γ -globin mRNA levels (Supplemental Fig. S1A), however at 24 hours a significant 1.4-fold (p=0.034) and 1.4-fold (p=0.032) increase in γ-globin by SAL and Hem (p<0.05) respectively was observed (Supplemental Fig. S1B). Furthermore, by Western blot analysis, SAL induced HbF at 6 hours 1.7-fold (p=0.041) and 2.2-fold (p=0.0078) by 24 hours (Supplemental Fig. S1C and S1D). These results demonstrate SAL induces HbF early time points but to see maximal effects we chose 48 hours based on previous *in vitro* experiments across all cells.

To further support a functional role of ATF4 in *HBG* regulation, we performed siATF4 gene silencing studies. K562 cells were transfected with scrambled siRNA (Scr) or 200nM siATF4 24 hours, followed by the addition of SAL (24μM) RNA for 48 hours. Using Scr as control normalized to one, siATF4 200nM silenced decreased gene by 50% (p=0.045) significantly (Fig, 2A). Next, we performed rescue experiments and showed SAL treatment overcame siATF4 inhibition and increase endogenous ATF4 levels by 3-fold (p=0.025). We also analyzed the effects on *HBG* levels under the same treatment conditions and showed siATF4 decreased *HBG* mRNA levels 47% (p=0.019), which was overcome by SAL treatment, activating *HBG* 2.1-fold (p=0.04) compared to siATF4 (Fig. 2B) However, SAL 24mM increased *HBG* by 2.5-fold above siATF4 treatment levels. We concluded SAL has the ability to overcome the silencing effect of siATF4 to activate *HBG* transcription supporting a role for ATF4 in *HBG* activation.

**Figure 2.**
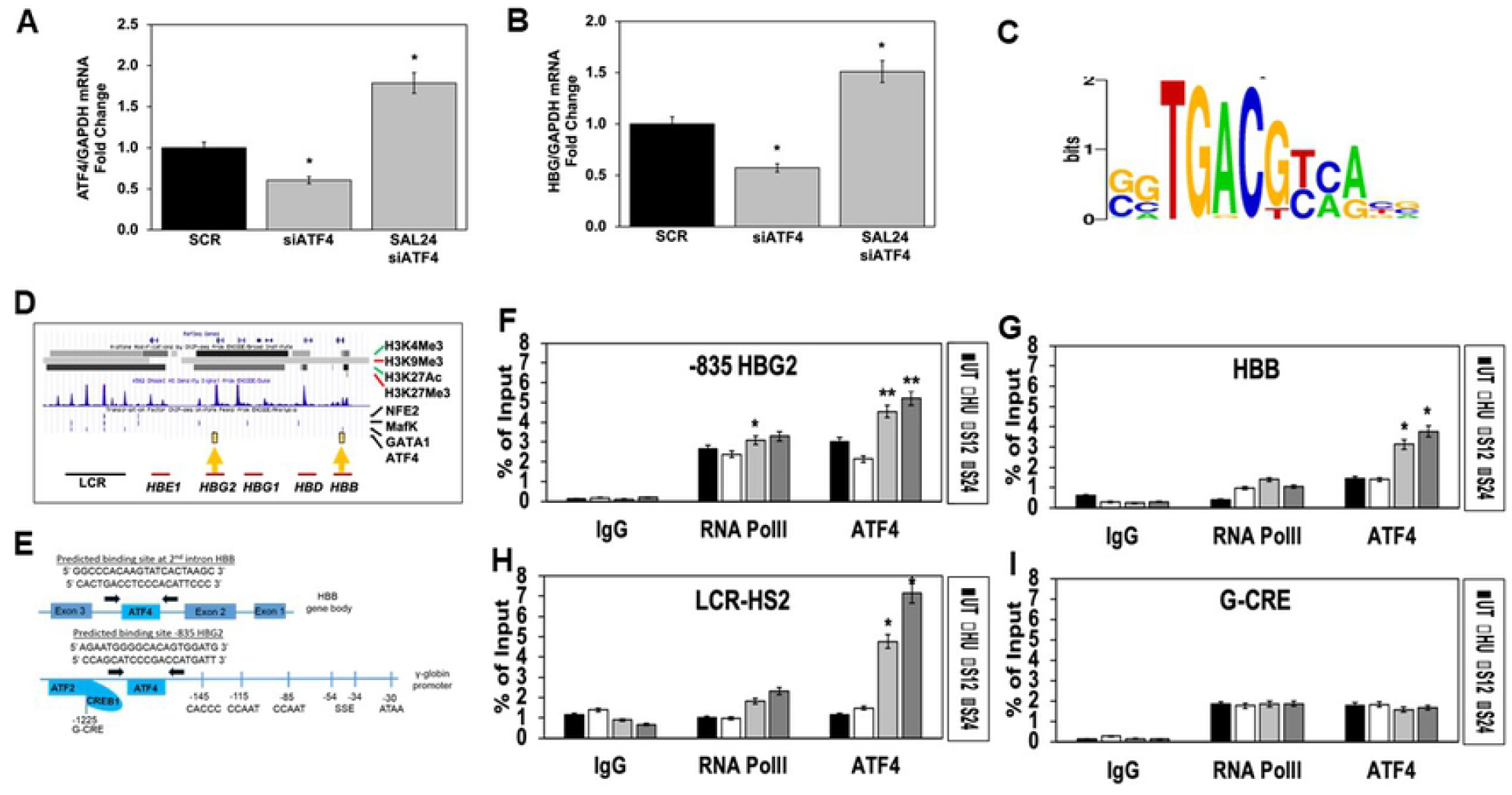
ATF4 binds to the *HBG* and *HBB* promoters to activate transcription. K562 cells were transfected with 200nM siATF4 or siRNA scrambled (SCR) control for 24 hours, followed by treatment with SAL (24μM) in triplicates for 48 hours. Total RNA was collected, and RT-qPCR analysis was performed to determine the levels of **A)** *ATF4* and **B)** *HBG* mRNA expression levels after siATF4 transfection. **C)** Integrated System for Motif Activity Response Analysis (ISMARA) software was used to generate the ATF4 consensus binding sequence: 5’GTGA CGT [A/C] [A/G]’3. **D)** ChIPseq for K562 cells was downloaded from the ENCODE database which showed strong ATF4 binding in the region of the predicted motif -835bp*HBG2* in the upstream promoter and weaker binding in the second intron of the *HBB* gene (yellow arrows). **E)** Schematic of ChIP assay primer sequences used for qPCR analysis. **F-I)** K562 cells were treated with HU (75μM; white bar), SAL 12μM (light grey) and SAL 24μM (dark grey) for 48 hours and used for ChIP assay using IgG (non-specific binding control), RNA polymerase II (PolII) (transcription rate control), and ATF4 binding at the **F)** -835HBG2, **G)** HBB gene, **H)** LCR-HS2 and **I)** G-CRE (n=3 per group). Data are shown as mean ± SEM and *p<0.05; **p<0.01; *** p<0.001 was considered statistically significant.

### SAL increase ATF4 binding in the *HBB* locus *in vivo*

To achieve *HBG* activation, we predicted ATF4 binding *in vivo* after SAL treatment, therefore potential ATF4 consensus motifs were identified using ISMARA (integrated system for motif activity response analysis) software. A consensus ATF4 (5’-TGACGTCA-3’) motif was identified in the G-CRE at -1225 upstream of *HBG2* (Fig. 2C). We previously demonstrated ATF2/CREB1 binding to the G-CRE that was required for HbF induction by sodium butyrate (17). Since ATF2 is a known binding partner of ATF4 (18), we examine ChIPseq data generated in K562 cells by the ENCODE project. We observed strong ATF4 binding at -835bp upstream of *HBG2* and a weaker binding site in the second intron *HBB* (Fig. 2D); this data was used to design qPCR primers (Fig. 2E).

To investigate *in vivo* DNA-protein interaction, ChIP assay was completed in K562 cells treated with SAL (12µM and 24µM) for ATF4 binding in the -835*HBG2* region, *HBB* gene, LCR-HS2 and G-CRE along with control RNA PolII (gene transcription rate) binding (Fig. 2F-2I). Since the *HBG* gene is actively transcribed in K562 cells, we observed chromatin enrichment for RNA PolII of 3.2-fold (p=0.015) and ATF4 up to 5.2-fold (p=0.011) at -835*HBG2* compared to IgG control (Fig. 2F). As expected, there was lower ATF4 binding at the *HBB* site (Fig. 2G). Since the LCR-HS2 is a powerful globin gene enhancer element, we investigated ATF4 binding in this region (19). Interestingly, there was a 7.2-fold (p=0.022) increase in ATF4 binding in the LCR-HS2 at SAL 24µM (Fig. 2H). The predicted ATF4 site in the G-CRE showed 2-fold chromatin enrichment however, which was not significantly changed by SAL treatment nor was RNA PolII binding (Fig. 2I).

### SAL activates *HBG* and *ATF4* transcription in human sickle erythroid progenitors

Hahn et al previously reported the ability of SAL to induce HbF in normal erythroid progenitors generated from CD34+ stem cells (2). Therefore, to provide additional support for developing SAL for treatment of SCD, we generated primary erythroid progenitors from PBMCs isolated from sickle cell patients as previously published (11). These cells are under oxidative stress and undergo hemolysis. After erythroid lineage commitment on day 8, we treated progenitors with SAL (9, 18 and 24µM) dissolved in water alone or combined with HU (75µM) for 48 hours. Cell viability remained >90% for all groups; RT-qPCR was performed and *HBG* and *HBB* mRNA were normalized by *GAPDH*. The calculated *HBG/HBB* mRNA ratios showed a significant 1.4-fold (p=0.004) and 1.6-fold (p=0.001) increase at SAL 18 and 24µm respectively (Fig. 3A), which was not increased further by combination SAL/HU treatment.

**Figure 3.**
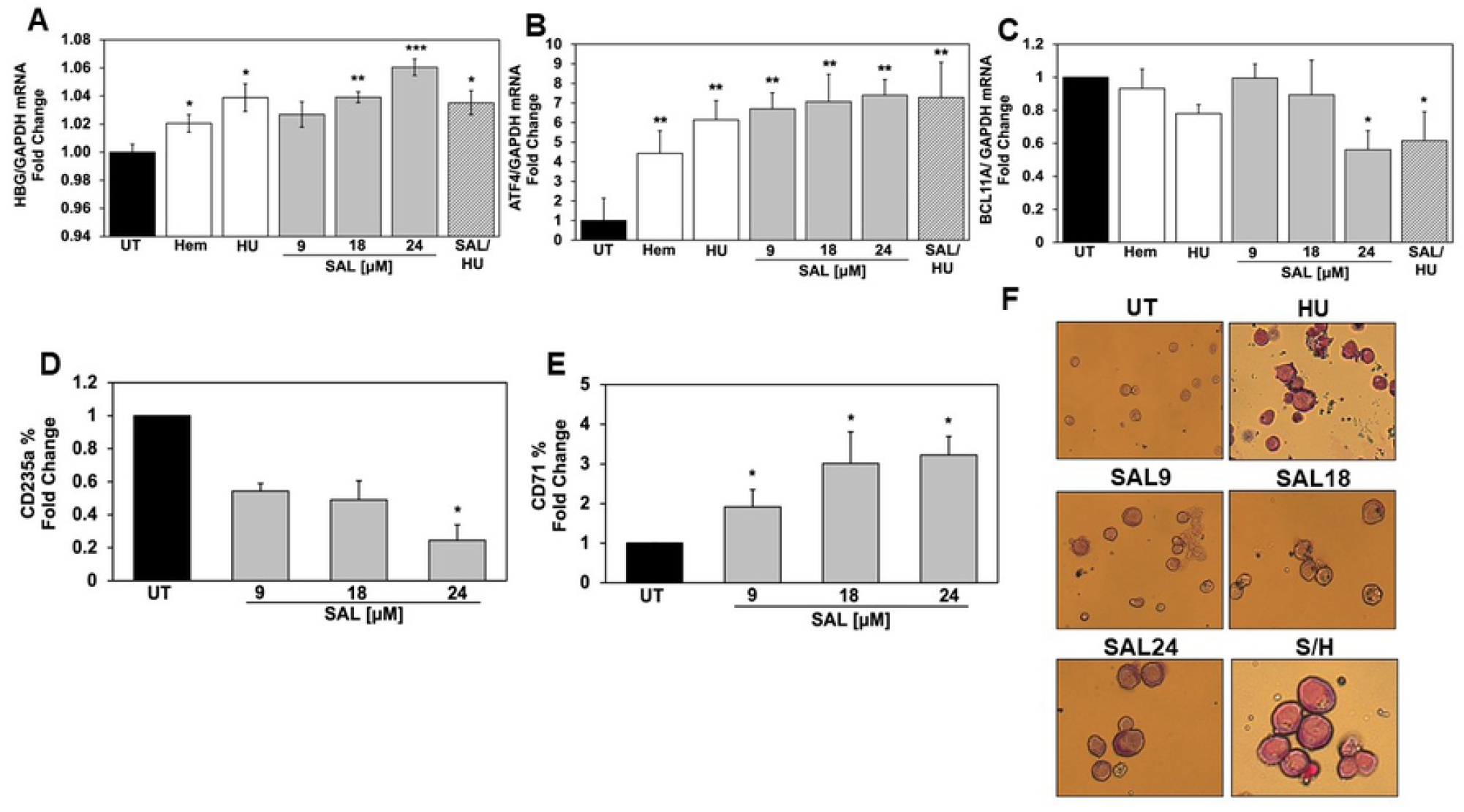
SAL activates HbF without changing HbS levels in sickle erythroid progenitors. Sickle erythroid progenitors were treated on day 8 with Hem 75μM, HU 75μM and SAL 9, 18 and 24μM alone or combined SAL 18μM combined with HU 75μM; SAL was dissolved in water. After 48 hours of treatment, data were generated and shown as the mean ± SEM (n=6) and *p<0.05; **p<0.01; *** p<0.001 was considered statistically significant. **A)** RT-qPCR demonstrated the *HBG/HBG + HBB* mRNA ratio calculated after the individual *HBG*, and *HBB* gene expression levels was normalized to *GAPDH* the internal control. RT-qPCR demonstrates **B)** *ATF4* mRNA levels and **C)** *BCL11A* mRNA levels normalized with *GAPDH* the internal control. Sickle erythroid progenitors under the different treatment conditions were stained with human FITC-conjugated **D)** CD71 and **E)** CD235a antibody and analyzed by flow cytometry. **F)** Representative images of erythroid progenitors generated at 40-x magnification by light microscopy.

To define molecular mechanism in sickle progenitors, we quantified ATF4 mRNA levels, which showed a significant dose-dependent increase of 6.7-fold (p=0.010), 7.1-fold (p=0.004) and 7.3-fold (p=0.0026) by SAL 9, 18 and 24µM, respectively (Fig. 3B). Interestingly, treatment with HU produced a 6.1-fold (p=0.025) increase in ATF4 suggesting an additive effect might not be expected for combination SAL/HU studies (Fig. 3A). We next tested the ability of SAL to silence the *HBG* repressor B-cell lymphoma/leukemia 11A (BCL11A). Similar to HU, SAL 24µM produced a 44% decrease in BCL11A (p=0.032) expression which was comparable to SAL/HU combination treatment (Fig. 3C).

To develop SAL for human trials, a better understanding of its effect on erythroid maturation is desirable. Therefore we measured transferrin receptor (CD71) and glycophorin A (CD235a), which are expressed on the surface of erythroid progenitors during differentiation (20). As shown in Fig. 3D, a dose-dependent 1.92-fold (p=0.030), 3.01-fold (p=0.037) and 3.2-fold (p=0.011) increase in CD71 expression was produced by SAL 9, 18 and 24µM, respectively. In addition, SAL mediated a dose-dependent decrease in CD235a expression by 44%, 51%, and 76% (p=0.027) at SAL 9, 18 and 24µM, respectively (Fig. 3E). Shown in Fig. 3F are Giemsa stains of progenitors under the different treatment conditions. These data suggest a dose-dependent delay in erythroid maturation by SAL.

### SAL induces HbF expression and prevents ROS in human sickle erythroid progenitors

Individuals with SCD experience oxidative stress produced by hemoglobin polymerization and chronic hemolysis with the release of free heme (21), therefore we investigated the ability of SAL to induce HbF and decrease ROS in sickle erythroid cells. On day 8, we treated progenitors with SAL (9, 18 and 24µM) or combined with HU (75µM) dissolved in water for 48 hours. Cell viability remained >90% for all groups. As shown in the representative histograms (Fig. 4A) and quantitative bar graph (Fig. 4B), treatment with SAL 9, 18 and 24µM increases F-cells from 15.3% (untreated) to 21.3%, 23.8% and 25.1% (p=0.021) respectively. Likewise, we observed an increase in HbF levels per cell using MFI (Fig. 4C) and Western blot analysis (Fig, 4D), reaching significance at the SAL 24µM (p=0.003) concentration; by contrast, HbS expression was not altered by SAL (Supplemental Figure S2 A-B). In the presence of SAL 24µM we observed a significant dose-dependent increase of HbF, p-eIF2α and ATF4 up to 1.6-fold (p=0.0441), 1.8-fold (p=0.043) and 1.5-fold (p=0.013) respectively (Supplemental Figure S2 C-E). Lastly, we determined changes in ROS levels, SAL treated cells were stained with DCF-DA (10 µM) for 4h and flow cytometry completed. Shown in Fig. 4E is representative histograms confirming a dose-dependent decrease in ROS by 30.8% (p=0.0000272), 63.8% (p=0.00000644) and 85% (p=0.000000133) with increasing SAL concentrations (Fig. 4F); adding HU did not improve ROS levels further.

**Figure 4.**
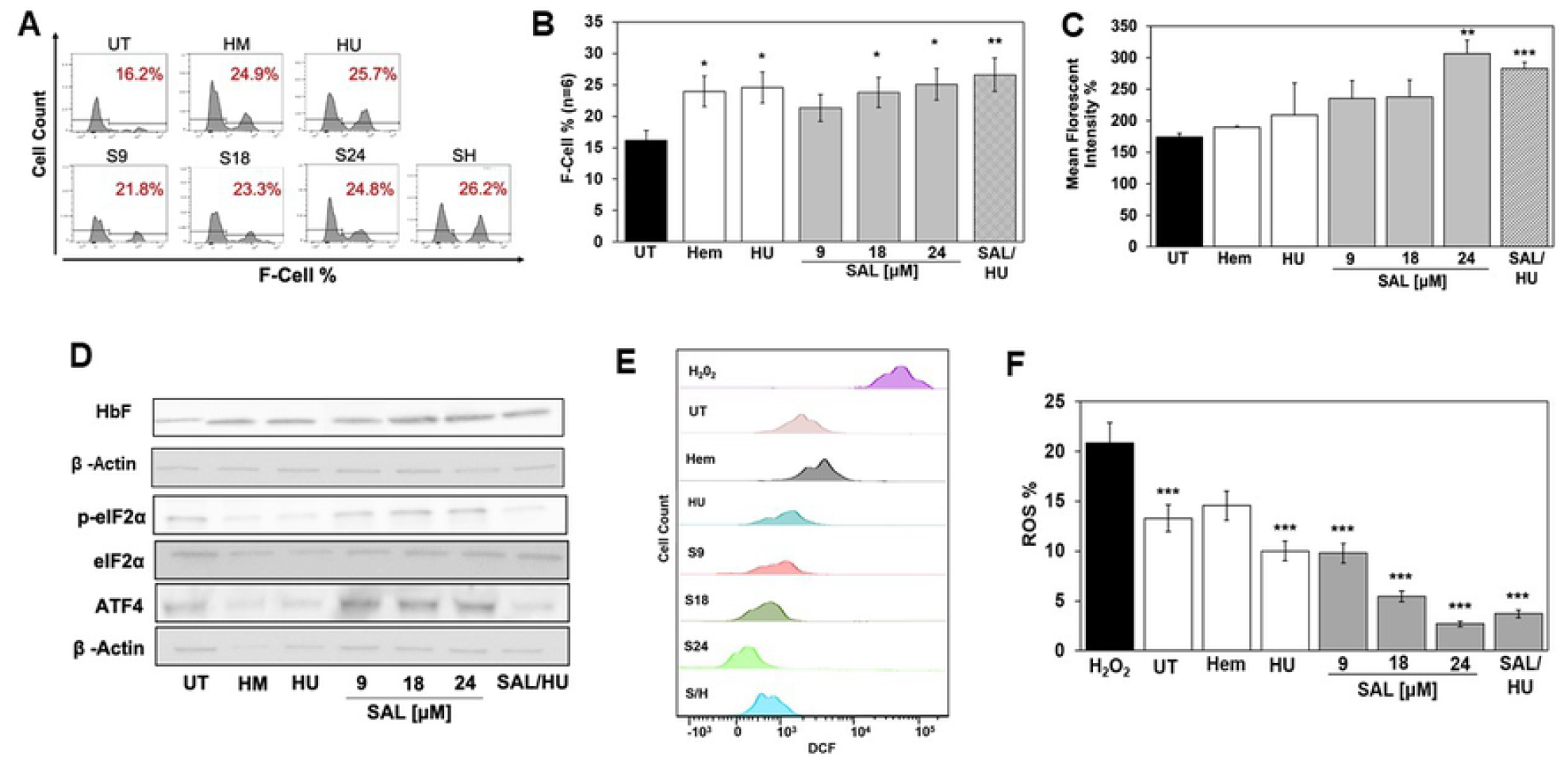
SAL activates HbF and reduces ROS levels in sickle erythroid progenitors. Sickle erythroid progenitors were grown in our two-phase culture systems and were treated on day 8 with Hem 75μM, HU 75μM and SAL 9, 18 and 24μM alone or SAL 18μM combined with HU 75μM; SAL was dissolved in water. After 48 hours of treatment, data were generated and shown as the mean ± SEM (n=6) and *p<0.05; **p<0.01; *** p<0.001 was considered statistically significant. **A)** Shown is histograms representation of F-cell levels determined by flow cytometry; **B)** quantitative data generated by FlowJo analysis. **C)** The level of HbF expression was measured by MFI data generated by flow cytometry analysis. **D)** Representative western blots gel of HbF, p-eIF2α and ATF4. **E)** Shown is a representative histogram overlay of DCF positive cell counts generated by flow cytometry as a measure of ROS levels; hydrogen peroxide (H_2_O_2_) was used as a positive control. **F)** Shown in the bar graph of the quantitative data of ROS levels compared to the untreated cells (UT).

### SAL induces HbF expression in SCD transgenic mice

Our long-term goal is to develop a novel agent for treatment of individuals with SCD; therefore, preclinical studies evaluating the pharmacokinetics characteristics, toxicity, and ability of SAL to induce HbF, were completed in Townes SCD transgenic mice (15). To establish the plasma distribution of SAL, liquid chromatography-multiple reaction monitoring mass spectrometry showed SAL parent ion peak at m/z 481.03 (Fig. 5A). Next, we treated mice with a single intraperitoneal (IP) SAL 5mg/kg dose and blood was drawn at 15 min, 30 min, and 1, 3, 6, 12 and 24 hours; SAL peaked at 6 hour (29 fmol) in plasma and decreased to barely detectable levels by 24 hours (Fig. 5B and 5C).

**Figure 5.**
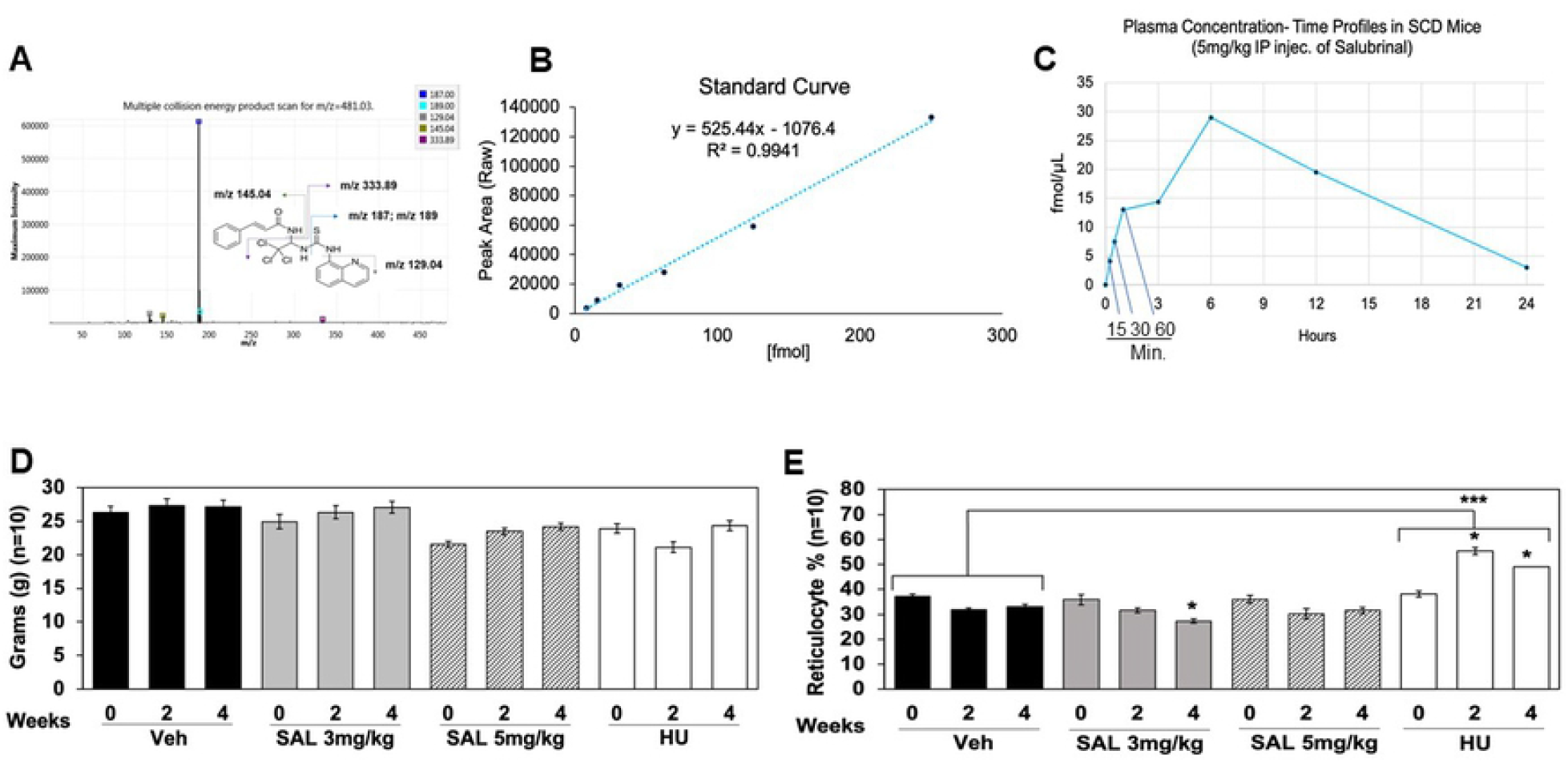
Plasma level of SAL in SCD transgenic mice and evaluation of toxicity. **A)** At the corresponding time points, blood was drawn by tail bleed into an EDTA tube. Plasma was immediately collected and 80µ l acetonitrile was added to 20µ l of mouse plasma and vortexed for 5 minutes. The samples were then centrifuged at 16,000g for 10 minutes and the supernatant was transferred into a glass vial for liquid chromatography-multiple reaction monitoring mass spectrometry. Multiple collision scan for the SAL molecule shows the different fragmentation by mass spectrometry. **B)** Shown is the standard curve used to determine plasma concentrations of SAL. **C)** The *in vivo* SAL levels were calculated from the standard curve (Panel B), using plasma isolated from peripheral blood of SCD mice. After establishing the plasma concentration of SAL, we next treated Townes SCD transgenic mice (4-6 months old) with SAL 3 mg/kg/day and SAL 5 mg/kg/day dissolved in water, for 4 weeks by intraperitoneal (IP) injections. Water vehicle control (Veh) and positive control hydroxyurea 100 mg/kg/day (HU) treatments were completed. All treatment groups consisted of n=10 mice with an equal sex distribution of 5 males and 5 females. **D)** Mouse weights in grams were obtained at week 0 (baseline), week 2, and week 4. **E)** Blood samples collected at weeks 0, 2 and 4 were stained with acridine orange for reticulocyte counts by flow cytometry. Data are shown as the mean ± SEM and brackets represent results of the ANOVA analysis across treatment groups. *p<0.05 and *** p<0.001 was considered significant.

To determine *in vivo* efficacy and toxicity, we treated SCD mice 4-6 months old with SAL 3 mg/kg/day (SAL3) and SAL 5 mg/kg/day (SAL5) or HU 100mg/kg/day (HU100) by IP injections, 5 days per week for 4 weeks (Supplemental Figure S4). Two independent replicates were completed with five mice each (n=10 per group). Animals were weighed and peripheral blood drawn at week 0, 2 and 4. Over 4 weeks of treatment, normal daily activity and weight gain was maintained (Fig. 5D) and no drug toxicity or significant change in blood counts was observed (Supplemental Figures S5 & S6). By contrast, HU produced a sustained significant decrease in granulocyte, monocyte and platelet counts by week 4. To evaluate the effect on erythropoiesis we measured reticulocyte count using acridine orange and flow cytometry analysis. Treatment with SAL3 produced a significant decrease in reticulocyte count from 35.9% to 27.3% (p=0.035) at week 4 which was not observed at the higher dose of SAL (Fig. 5E), while treatment with HU100 increased reticulocytes up to 49.15% (p=0.033). To determine whether SAL and HU treatment changed reticulocyte counts significantly compared to vehicle control, ANOVA was performed with n=10 mice per group. Of note, the minor decrease at SAL3 week 4 was not different from control. By contrast, HU100 significantly increased reticulocytes counts (p=0.00018) compared to water control (Fig. 5E, brackets). While significant changes in hemoglobin or hematocrit were not mediated by SAL, more importantly, there was no signs of toxicity on hematopoiesis. The variability of individual response to treatment recapitulates patterns observed in human clinical trials for HU in sickle cell patients.

To establish whether SAL mediated *in vivo* HbF induction, we performed flow cytometry analysis. For SAL3, F-cells increased from 3.7% to 8.5% (p=0.0013) and SAL5 increased F-cells to 8.4% (p=0.0023) (Fig. 6A). This level of F-cell induction was comparable to the 8.4% increase for HU100 (p=0.014); shown in Fig. 6B, is representative histograms of F-cells data. ANOVA analysis demonstrated significant differences in F-cells for mice treated with SAL3 (p=0.0357), SAL5 (p=0.0421) and HU100 (p=0.0305) compared to water vehicle control. We also quantified HbF levels per cell by MFI analysis, where we observed a significant increase from 297 to 542 units (p=0.00014) for SAL3 and SAL5 changed from 335 to 502 units (p=0.0044) (Fig. 6C). Similarly, HU increased MFI to comparable levels suggesting SAL might be effective *in vivo* similar to HU in SCD. To further support SAL clinical development, we tested its ability to achieve an anti-sickling effect. Mouse blood was examined by light microscopy (Fig. 6D) and quantitative data confirmed SAL5 reduced the percentage of sickled erythrocytes by 66% (p=0.00319) (Fig, 6E). These findings support the ability of SAL to induce HbF and produce phenotypic changes in the preclinical SCD transgenic mouse model, similar to HU without significant side effects or toxicity.

**Figure 6.**
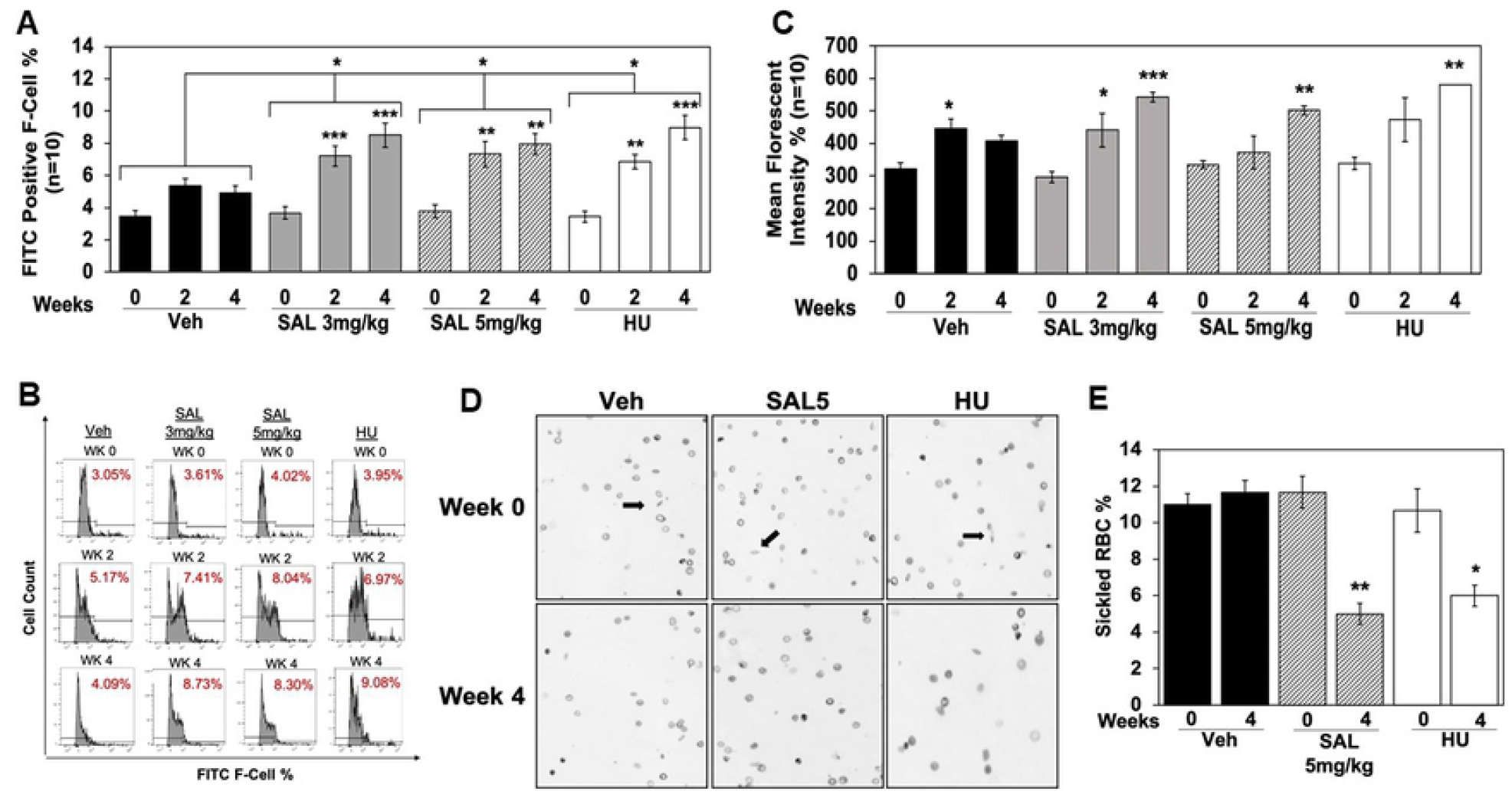
SAL increases HbF expression and produce anti-sickling effects in SCD mice. **A)** Peripheral blood collected at the time points shown was stained with FITC-anti-HbF antibody and flow cytometry performed to quantify F-cells by FlowJo analysis. The paired student’s t-tests were used to determine the significance in F-cells at week 0 compared to week 2 and 4 within each group (n=10). ANOVA was used to compare the significance of F-cells between water compared to the two concentrations of SAL and HU group; results are represented by brackets between treatment group. Data are shown as the mean ± SEM and *p<0.05; **p<0.01; *** p<0.001 was considered significant. **B)** Representative histograms demonstrating the increase in the number of FITC-HbF positive cells from week 0 to week 4 for each treatment group. **C)** The level of HbF expression per cell was measured by MFI data generated by flow cytometry analysis. **D)** Shown is a representative image generated at 40-x magnification by light microscopy. **E)** The levels of sickled red blood cells for the different conditions (black arrows) were quantified for 500 cells/triplicate and shown in the graph.

## DISCUSSION

Identifying molecular mechanisms involved in *HBG* activation will support development of additional novel therapies for individuals living with SCD. Hydroxyurea is the only FDA-approved HbF inducing drug used in SCD which ameliorates the clinical phenotype of frequent pain episodes, inflammation and hemolysis, and improves long term survival (22). Other potent HbF inducer tested in clinical trials such as arginine butyrate (23, 24), 5-azacytidine (25), decitabine (26) and short chain fatty acid derivatives (27)) function through diverse molecular mechanisms but none have advanced to FDA-approval for SCD. Factors such as a requirement for intravenous administration of arginine butyrate, hindered development since the agent is rapidly inactive when given orally (28). Likewise, decitabine mediated HbF induction in a Phase 1 clinical trial when given intravenously, however lack of efficacy by oral administration required combination treatments with tetrahydouridine, which has shown promise in clinical trials with SCD patients (29). Therefore, the discovery of non-chemotherapeutic oral HbF inducers that act by unique molecular and cellular mechanisms for use as a standalone agent or combined with HU support development of additional safe and effective treatment options for SCD.

We investigated the ability of the SAL to induce HbF expression through the eIF2α-ATF4 stress-signaling pathway in normal and sickle erythroid progenitors. SAL is a semi-permeable synthetic compound and is a selective inhibitor of PP1, interfering with the recruitment of p-eIF2α to PP1 through GADD34 and CReP. Under stress conditions, p-eIF2α rapidly inhibits general protein synthesis, including globin chains to prevent cell toxicity. Moreover, p-eIF2α increases translation of selective mRNAs, such as ATF4 to reprogram gene expression for adaptation to stress.

Severe cellular oxidative stress associated with chronic hemolytic states is predicted to promote cell death via ATF4-mediated activation of C/EBP homologous protein (30). Previously our group established the ATF2/CREB heterodimer binds the consensus G-CRE motif to induce HbF expression in primary erythroid cells (31). Since ATF4 binds in the G-CRE, we investigated p-eIF2α-ATF4 stress signaling as a mechanism involved in HbF induction by SAL. To support this speculation, we observed increases in p-eIF2α and ATF4 levels and *in vivo* binding of ATF4 at the -835*HBG* and *HBB* predicted motifs by ChIP assay, replicating published ENCODE ChIPseq data. In addition, there was an increase of ATF4 binding at the LCR-HS2 enhancer, which is required for DNA loop formation to activate globin gene transcription during hemoglobin switching. The weaker *HBB* motif showed low ATF4 binding however, we observed not increase in HbA or HbS *in vitro*. We speculate since the motif is not a promoter or enhancer element along with weaker histone marks that ATF4 did not support activation of the *HBB* gene.

In all cells tested, SAL mediated a dose-dependent increase of F-cells and HbF expression with a parallel decrease in ROS levels, thus supporting dual beneficial effects of treatment with this agent. In fact, treatment with hemin recapitulates the high oxidative stress produced by free heme; this mechanism for ATF4 activation as a stress response supports a relationship between hemin and ATF4 (32). To explicate the role of ATF4 in *HBG* activation, siATF4 studies showed the ability of SAL treatment to overcome gene silencing and rescue *HBG* activation. To evaluate the effects on erythropoiesis, we observed SAL 24µM treatment increased CD71 and decreased CD235a cell surface expression suggesting delay in erythroid progenitor maturation (33). Interestingly, in our SCD mouse studies the higher SAL dose produced a slight increase in reticulocytes, which correlates with CD71^+^ cells therefore, what was observed in tissue culture did not recapitulate in mice where there were no adverse effects on peripheral blood counts.

SCD pathophysiology includes chronic hemolysis, anemia and organ damage caused by HbS polymerization under deoxygenated conditions; as a result, the lifespan of red blood cells is shortened to 14-21 days due to oxidative membrane damage (34). It is known that chronic hemolysis leads to excess hemoglobin in plasma and oxidative stress complicated by free heme, haptoglobin deficiency and globin chain precipitation (35). Since we observed decreased ROS levels with SAL treatment there are added benefits of protecting SCD progenitors against oxidative stress. In addition, SAL produced phenotypic antisickling effects in SCD mouse blood, which is clinically relevant.

The dose and duration of SAL is critical in achieving its efficacy for enhancing HbF production. We are the first to utilize sickle erythroid progenitors generated from PBMCs from humans to test the ability of SAL to induce HbF expression. To guide our dosing, Suragani et al., suggested that higher concentrations than 25µM SAL would further increase p-eIF2α resulting in inhibition of protein synthesis (32). Other *in vitro* studies have shown SAL 3-24 µM activates *HBG* transcription in normal erythroid cells (2).

Heme-regulated eIF2α kinase activates the p-eIF2α-ATF4 integrated stress response in erythroid precursors to control ROS levels. In iron deficiency states, signaling through this pathway regulates globin chain synthesis and reduce oxidative stress during erythroid differentiation (32). Increases in p-eIF2α inhibit globin mRNA translation to prevent chain imbalance due to limited availability of heme. Furthermore, p-eIF2α enhancement increase protein chaperone reserves in cells, to aide in protein folding during stress (36). We showed that SAL changes the *HBG/HBB* mRNA ratio at steady state, along with *ATF4* expression.

A recent study confirms HRI-eIF2aP-ATF4 signaling suppressed *HBG* transcription by enhancing *BCL11A* expression (37), however, this mechanism has not been studied in sickled erythroid progenitors under oxidative stress. Blobel, et al. (37) demonstrated that ATF4 activates *BCL11A* in HUDEP2 cells, however, the mechanism did not show the same effect in their HRI^™/™^ mouse model, concluding that this mechanism can vary between cells. By contrast, in our sickle erythroid progenitor studies, SAL activated ATF4 expression and decreased *BCL11A*, similar to HU. Furthermore, treatment with siATF4 decreased *HBG* mRNA in K562 cells, which were rescued by SAL supporting a role for ATF4 in SAL mediated HbF induction.

Studies have shown that ATF4 regulates a variety of genes that control cell adaptation to stress conditions, however, long-term stress results in C/EBP homologous protein activation and initiation of apoptosis (38). SAL inhibits the negative feedback loop of the integrated stress response pathway to promote increased p-eIF2α levels and ATF4 activation. Furthermore, limited data exist to demonstrate a role of ATF4 signaling in erythroid differentiation however, our data support the ability of SAL to activate ATF4 and reduce oxidative stress in SCD progenitors. In addition, the pathogenesis of neurodegeneration in brain involves ATF4 expression. Wu et al. (39) showed that treatment of mice with SAL up-regulated ATF4 expression. Subsequent studies with siATF4 showed silencing of the parkin protein required for protection against rotenone cytotoxicity. These data indicate the ATF4-parkin pathway plays an important role in the SAL-mediated neuroprotection of rotenone-induced dopaminergic cell death.

To provide further insight and evidence of efficacy for the development of SAL as a non-chemotherapeutic HbF inducer for individuals with SCD, preclinical studies were completed in the Townes SCD mouse model which has been used to test various agents for their capacity to induce HbF *in vivo* (40, 41). Chronic treatment demonstrated the ability of SAL to induce HbF and produce an anti-sickling effect without toxicity. We know SCD patients experience endoplasmic reticulum stress due to mast cell activation, mitochondrial dysfunction, and associated oxidative stress to activate the integrated stress pathway (42). Gupta and colleagues (42) demonstrated the ability of SAL to attenuate pain in the transgenic HbSS-Berk mouse model, with concomitant decrease in ROS and endoplasmic reticulum stress (43).

Collective data generated in two SCD mouse models support development of SAL as a novel agent for treatment of SCD. Our studies show an innovative and novel approach using the sickled erythroid progenitors from patients and the Townes sickle mice showing consistent *in vitro* and *in vivo* data. In addition to SAL increasing HbF and decreasing ROS production, red blood cell sickling is reduced. These data support inhibition of vaso-occlusive pain episodes demonstrated by Gupta and colleagues (43). Although we did not observe an additive effect of SAL with HU, SAL performed at a comparable level to HU as it relates to HbF induction and reducing oxidative stress.

SAL is under development for other diseases such as hepatic steatosis by altering cellular stress and autophagy through eIF2α signaling (44), treatment of osteogenesis imperfect (45), and SAL efficiently blocked osteoporosis in mice (46). To move SAL to clinical trials an oral formulation is required; mice have been treated with SAL in soymilk, which restored primordial follicles production in galactosemia mice (47) however, the design of novel and safer alternatives for HU is highly desirable for people with SCD.

## ACKNOWLEDGEMENT

We would like to thank Ms. Krystle Stone, research assistant in the Pediatric Sickle Cell program for faithfully collecting discard blood samples from patients on chronic transfusions, which was used to isolate peripheral blood mononuclear cells. We would also like to thank Dr. Xingguo Zhu, for his expertise on the protein studies and assistance with the mouse studies.

## FUNDING

This work was funded through grants from the National Heart Lung and Blood Institute, HL069234 to BSP and NL (Diversity Supplement); and SBIR/STTR, R42 HL136068 to BSP and NL (Diversity Supplement).

### Author contributions

Sanam Loghavi and Sa A. Wang designed the study; Sanam Loghavi, Ken Furudate, Tomoyuki Tanaka, Noushin R. Farnoud, Rashmi Kanagal-Shamanna, Sherry Pierce, Keyur P. Patel, Joseph D. Khoury, Jeffrey L. Jorgensen and Sa A. Wang collected and analysed data; Courtney DiNardo, Koichi Takahashi, Nicholas J. Short, Tapan Kadia; Marina Konopleva, Naval Daver, Musa Yilmaz, Hagop Kantarjian, Farhad Ravandi enrolled patients; Sanam Loghavi, Sa A. Wang, Courtney DiNardo, Marina Konopleva, L. Jeffrey Medeiros wrote the manuscript; all authors have read and approved the final copy of the manuscript

## AUTHOR CONTRIBUTIONS

NHL contributed to experimental design, performed experiments, collected and analyzed data and wrote the manuscript; BL contributed to primary culture studies and drug treatment design; CP contributed to ChIP Assay, siRNA studies; UN contributed to mouse genotyping by HPLC and reviewed paper; MZ established the Townes transgenic mouse and contributed to mouse studies; HX created database and performed statistical data analysis for tissue culture and mouse treatments; WZ contributed to pharmacokinetic studies and mass spectrometry; BSP conceived and designed the study and revised the manuscript. Authors approve the final version of manuscript.

